# scRegulocity: Detection of local RNA velocity patterns in embeddings of single cell RNA-Seq data

**DOI:** 10.1101/2021.06.01.446674

**Authors:** Akdes Serin Harmanci, Arif O Harmanci, Xiaobo Zhou, Benjamin Deneen, Ganesh Rao, Tiemo Klisch, Akash Patel

**Author notes:** These authors contributed equally (Akdes Serin Harmanci, Arif O Harmanci).

## Abstract

Single cell RNA-sequencing has revolutionized transcriptome analysis. ScRNA-seq provides a massive resource for studying biological phenomena at single cell level. One of the most important applications of scRNA-seq is the inference of dynamic cell states through modeling of transcriptional dynamics. Understanding the full transcriptional dynamics using the concept named RNA Velocity enables us to identify cell states, regimes of regulatory changes in cell states, and putative drivers within these states. We present scRegulocity that integrates RNA-velocity estimates with locality information from cell embedding coordinates. scRegulocity focuses on velocity switching patterns, local patterns where velocity of nearby cells change abruptly. These different transcriptional dynamics patterns can be indicative of transitioning cell states. scRegulocity annotates these patterns with genes and enriched pathways and also analyzes and visualizes the velocity switching patterns at the regulatory network level. scRegulocity also combines velocity estimation, pattern detection and visualization steps.

## Introduction

Single-cell RNA Sequencing has enabled us to study heterogeneous cell populations with single cell resolution. Technologies such as 10X Genomics Chromium^1^, inDrop^2^, SMART-seq2^3^, Drop-Seq^4^ are used to sequence transcripts from thousands of cells that are isolated from a sample of interest. Current data analysis pipelines analyzing scRNA-Seq data reveals a static snapshot of cellular states. Standard scRNA-seq pipelines focus on quality control^5–9^, cell filtering ^5–9^, dimensionality reduction ^5–9^, integration^10–17^, differential expression^18–20^, and clustering of the cells^5,8,9^, and assignment of cell types from the samples^21–25^. These are very important steps to organize and assess the quality of the single cell datasets and provide an initial analysis of the data. As single cell datasets tend to be very large with possibly hundreds of thousands of cells, these initial analyses provide important insight into the biological states of the cells and the studied conditions. As such, the single cell datasets contain massive amount of information that should be extracted with computational and statistical methods. One of the main challenges is that datasets are being generated at a rate that is much faster than the computational methods can analyze.

Although scRNA-seq is generated from a single time point, it can be used to estimate the transcriptional dynamics of transcriptional states of cells to study, for example, developmental, disease, and immunological processes that exhibit large dynamic changes in cells^26–29^. Understanding transcriptional states enables us to define cell types and cell-type defining markers more coherently. Additionally, it allows us to infer the heterogeneity of tumor samples at the single cell level.

One way to infer transcriptional dynamics is through trajectory analysis. The main hypothesis for these analyses is that the sample comprise heterogeneous sets of cells from a continuum of dynamic states. These states can represent dynamic processes such as differentiation and cell cycle^30–35^. These states can be modeled by numerous “trajectories” where the dynamic states are connected on a trajectory of states (e.g., Markov processes). Methods that perform trajectory analysis assume that the continuum of cellular states are sufficiently observable in the single cell RNA-seq sample. The idea is to build parsimonious trajectories that explain the changes in the cell types. The trajectory analysis is often coupled with pseudotime analysis^36,37^ to assign relative time units to the dynamic trajectory of the cells. This way, the cells in each of the trajectories can be aligned properly.

There are also methods that extract dynamicity information from all the cells at the same time by estimating the derivative of gene expression levels. This is performed by concept named RNA velocity, that can reliably estimate the relative time derivative of the gene expression state. RNA velocity enables us to study cellular transcription kinetics using the ratio of spliced and unspliced read counts of each gene across RNA-Seq data ^28,29,38,39^. The underlying assumption in this model is that genes are initially transcribed in an unspliced manner and then spliced, such that observed intronic reads can be interpreted as corresponding to nascently transcribed mRNAs. Transcriptional upregulation of a gene will result in a transient excess of nascent (unspliced) transcripts compared with processed (spliced) transcripts, whereas transcriptional downregulation results in a relative depletion of nascent (unspliced) transcripts.

Unlike trajectory analysis, RNA-velocity does not require the cellular composition to be diverse enough since each cell is processed by itself and velocity can, in principle, be estimated in each cell independently. Although several methods have utilized RNA-velocity for building trajectories and estimating cellular dynamics at a sample-wide (or global) level, there is still much information to be extracted from “local patterns”, i.e. the interactions of subsets of cells have with each other. The “local patterns” can be more concretely described by considering embeddings of cells in lower dimensions where the nearby cells in the embeddings are more similar to each other in terms of transcriptional states. One example of these is tSNE and UMAP based embeddings that are used extensively for visualizing scRNA-seq datasets. The local patterns in the embeddings can provide an incredible amount of biological insight. The relationship between velocity and the cellular dynamicity at the level of localities in the embeddings is not well-studied. We hypothesize that there is a need to develop new computational methods for detecting, summarizing, and visualizing the biological insights of dynamicity of cellular states in connection to the embeddings.

In this study, we present the scRegulocity algorithm, for measuring the dynamics of gene expression in large numbers of single cells using RNA velocity. Specifically, scRegulocity detects genes with velocity switches whereby the gene exhibits a strong velocity difference among cells that are nearby in the coordinates of embeddings. The local velocity switching patterns are very frequently observed in manual inspection of the estimated velocity distributions on the cells. We believe that the genes with velocity switches potentially represent drivers of dynamic processes such as cellular differentiation/development and disease progression, and these can be instrumental to delineate the drivers of dynamicity of these processes especially in tumors and cancers. In particular, these genes exhibit transcriptional dynamics which are detected by integrating RNA velocity and expression (to build the embeddings) rather than using expression signatures alone. ScRegulocity also reconstructs gene regulatory networks in transitioning cell states using RNA velocity. Our analyses on different single cell RNA-Seq datasets show that scRegulocity can recover driver TFs and transcriptional programmes in transitoning cell states which can not be easily inferred from whole transcriptome data. We believe that scRegulocity will facilitate the study of gene regulation in diverse biological systems.

Compared to other methods, scRegulocity stands out as a “local dynamicity inference” tool, whereas the majority of the other tools aim at detecting and describing patterns at a sample-wide level (or globally). scRegulocity takes standard files as inputs, is flexible and can be integrated into scRNA-seq analysis pipelines.

## Results

### scRegulocity Algorithm

Figure 1 illustrates the scRegulocity algorithm. The input is the aligned RNA-seq reads (e.g., SAM/BAM file) and the list of cell ids that will be analyzed. scRegulocity provides an integrated and complete pipeline starting from mapped reads and uses a spatial signal processing approach to detect the velocity switching patterns on embeddings. A velocity switching pattern is defined by an abrupt coordinated increase (or decrease) in velocity between two groups of cells that are close in the embeddings. Thus, it is vital for the embedding of cells into lower dimensions to provide useful biological information for nearby cells that are close to each other. For most of the embeddings that are widely used (such as tSNE, UMAP, and PCA) closeness generally implies biological similarity and therefore should be meaningfully usable in the context of velocity-switching analysis. In this study, we focus on tSNE and UMAP-based dimensionality reduction using the gene expression counts, i.e., the embeddings represent similarities in the global transcriptomic profiles. The velocity switching patterns are expected to identify the cells that are similar in transcriptional state but harbor opposite dynamic changes in expressional states that potentially stem from the regulatory state of the cells. One of the motivations for developing scRegulocity is that the velocity switching patterns are frequently observed in manual inspection of the expression velocities after they are mapped on the embedding coordinates.

**Figure 1.**
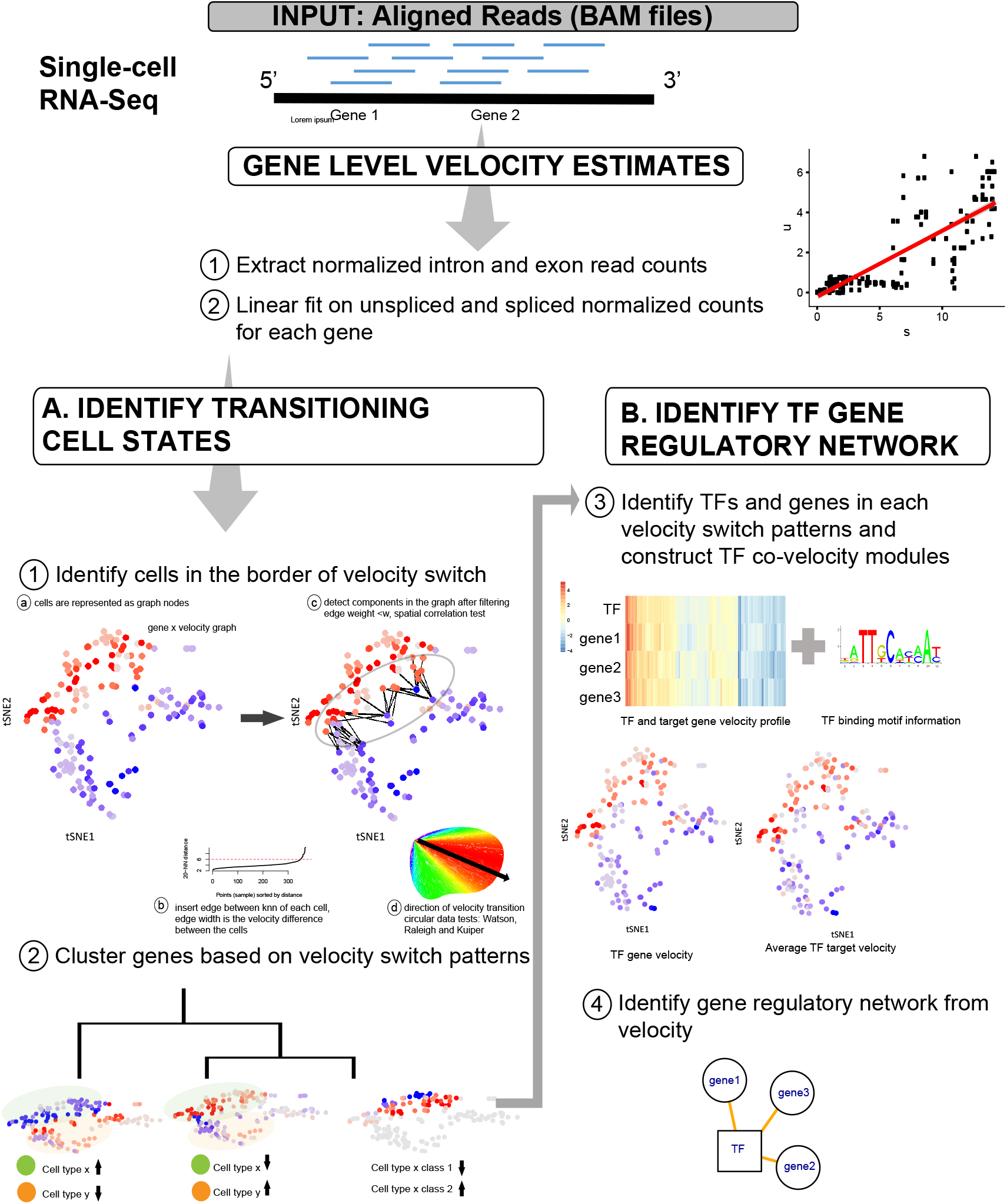
Overview of scRegulocity algorithm

In order to comprehensively characterize the velocity-switching patterns at the regulatory level, we mapped known TF-target interactions onto the genes that exhibit significant velocity switching-patterns. Then scRegulocity classifies the concordance or discordance of velocity switches with regulatory relations between the TF and target at the velocity-level and/or expression-level. Finally, the identified regulatory switches and regulatory information are visualized on the embeddings. scRegulocity can generate the popular embeddings of the cells after quantifying the expression levels on each cell. However, the user can skip this step if he/she has already generated the embedding themselves. We describe the other steps of scReguloCity workflow in Methods section.

### Accuracy of velocity estimates using sci-fate data

We first applied scRegulocity algorithm on cortisol response dataset generated from a method named sci-fate^40^. Sci-fate method is a combined single-cell combinatorial indexing and mRNA labelling to profile the ‘older’ and ‘newer’ transcripts based on their splicing status in single cell resolution. In this study, researchers identified regulatory elements and transcriptional drivers in cortisol response using the newly synthesized expression values. The newly synthesized expression values in scifate-study is the ground truth for the RNA Velocity values. Therefore we validated our scRegulocity algorithm using the cortisol response dataset and detected similar transcriptional programs reported in the scifate study.

In order to show the similarity of velocity and newly synthesized expression, we first calculated the correlation of velocity with newly synthesized and whole-transcriptome data for each gene separately. Figure 2A shows the distribution of correlation for each gene. We have detected a higher mean correlation with velocity and newly synthesized data compared to whole-transcriptome data (Figure 2A). We also checked the correlation of velocity of TF with TF target genes reported in sci-fate study (Figure 2B). We observed a higher correlation between velocity of TF with its target gene velocity values. Thus, we can infer the TF target regulatory network more accurately using RNA velocity values.

**Figure 2.**
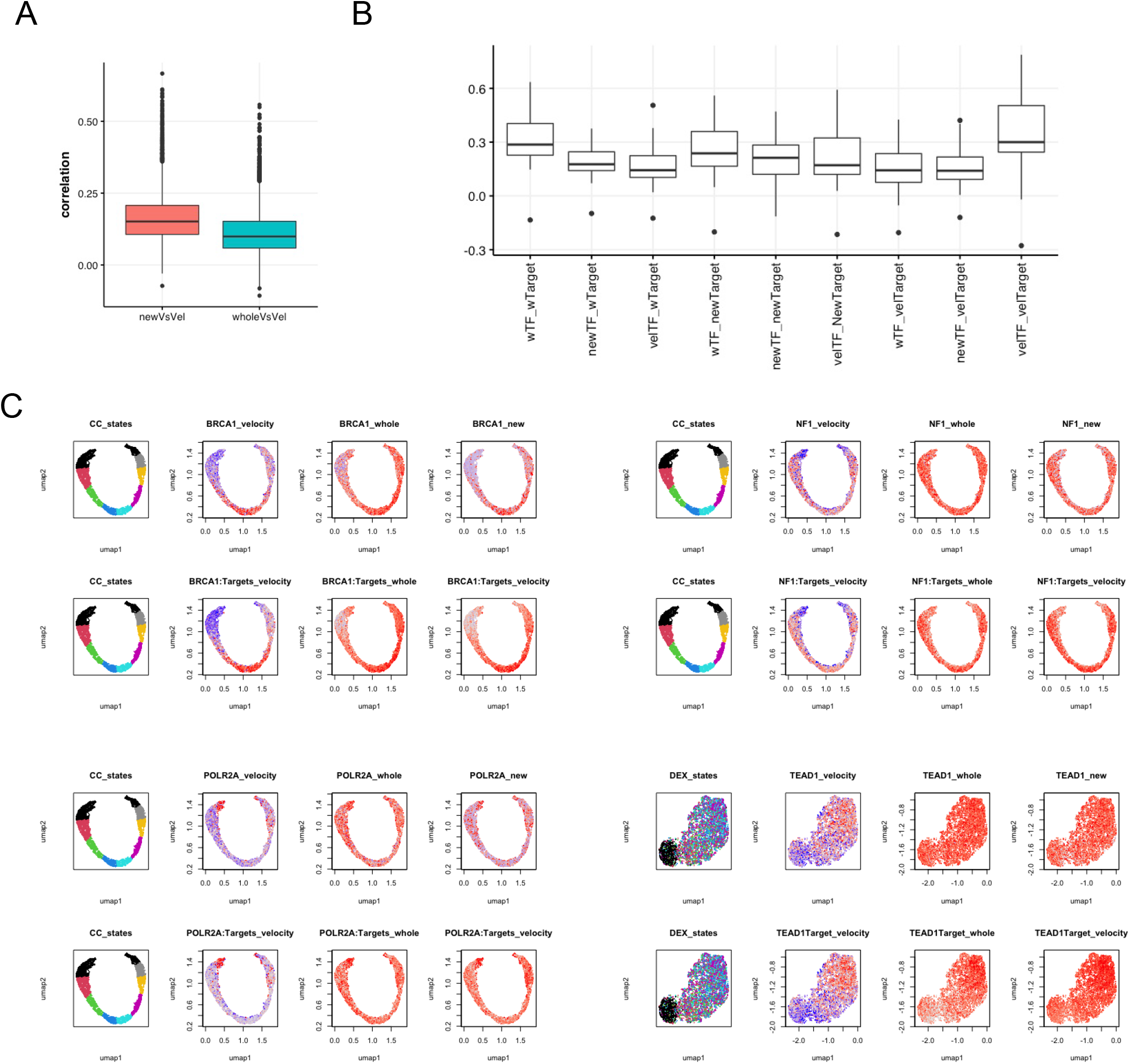
**A)** Correlation of velocity with newly synthesized and whole-transcriptome data for each gene separately **B)** Correlation of velocity of TF with TF target genes reported in sci-fate study **C)** RNA Velocity of cell cycle TFs such as *POLR2A, NF1, BRCA1* and GR response TFs such as *TEAD1* were highly correlated with the levels of velocity.

We next identified genes that have uniform direction of the velocity switch vectors in a subset of cells in order to define driver genes in cell state transitions. The velocity of cell cycle TFs such as POLR2A, NF1, BRCA1 and GR response TFs such as TEAD1 were highly correlated with the levels of velocity, more so than overall target gene mRNAs (Figure 2C).

### scRNA-seq of Chromaffin differentiation

We next applied our scRegulocity algorithm on a Chromaffin differentiation dataset studied in velocyto paper^41^. Researchers detected a movement of the differentiating cells towards a chromaffin fate using RNA Velocity. We first detected genes with velocity switches among different cell states and types. ScRegulocity identified Serpine2 having a significant velocity switching pattern among SCP (schwann cell precursor) cells and Differentiation cells (Figure 3A). We next clustered genes using velocity values and identified different velocity switching patterns (Figure 3B). Then we performed enrichment analysis on the genes within each cluster. We next sought the TFs that drive the progression of chromaffin fate differentiation, and inferred TF target regulatory network using RNA velocity values (Figure 3C). Chromaffin differentiation related TFs, such as Gata3, Tcf7l2, Sox6 and Tcf4, were identified using scRegulocity and the TF velocity values of these TFs were highly correlated with the mean TF target velocity values (Figure 3D).

**Figure 3.**
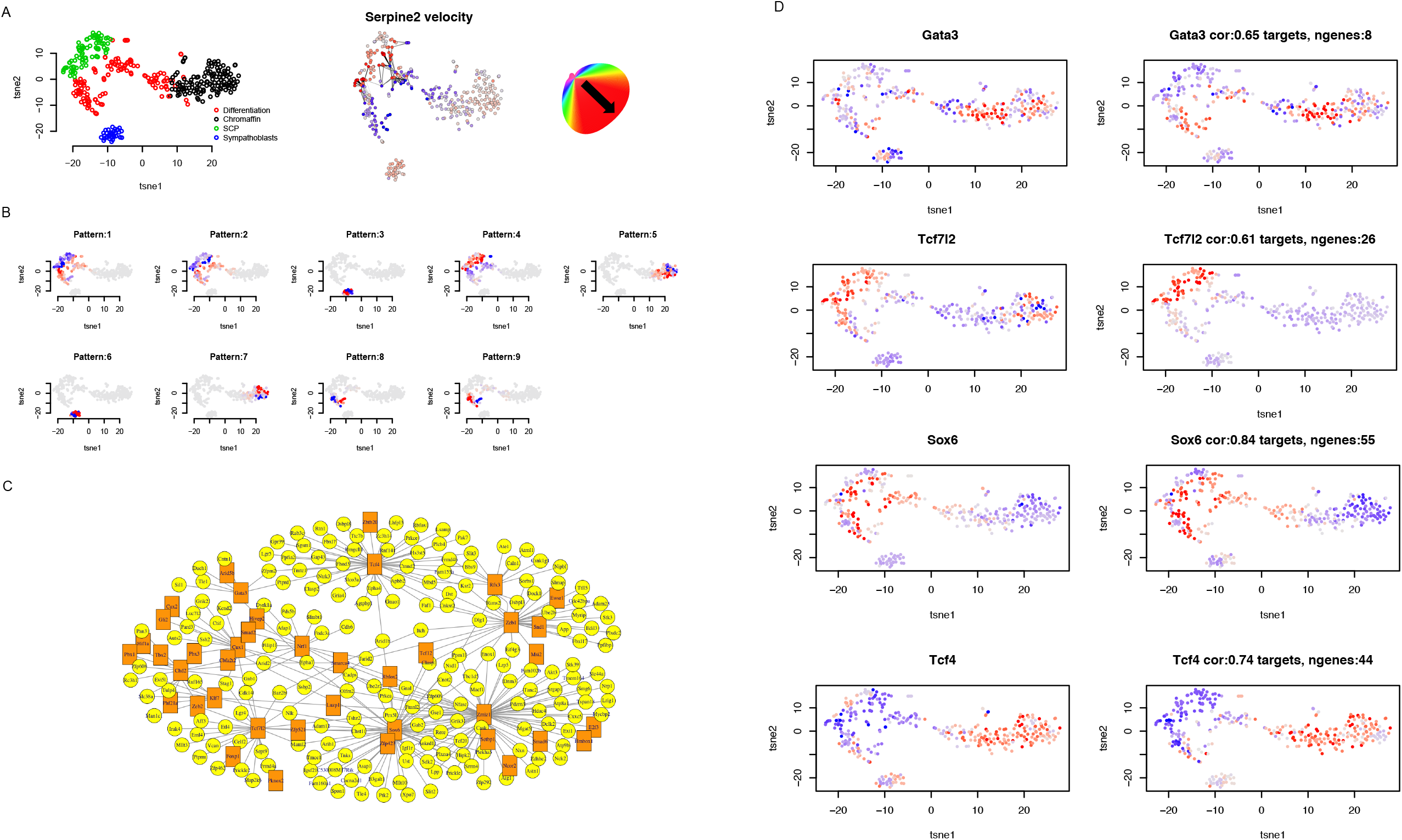
**A)** ScRegulocity identified Serpine2 having a significant velocity switching pattern among SCP (schwann cell precursor) cells and Differentiation cells **B)** velocity switching patterns identified by ScRegulocity **C)** identified TF target regulatory network using RNA velocity values **D)** Chromaffin differentiation related TFs, such as *Gata3, Tcf7l2, Sox6* and *Tcf4*, were identified using scRegulocity and the TF velocity values of these TFs were highly correlated with the mean TF target velocity values.

### Single-cell RNASeq Meningioma

We also applied our algorithm on our scRNA-Seq meningioma dataset. In total, after filtering low quality cells we have n=12244 cells from n=2 NF2 mutant recurrent meningioma tumors. We first clustered our scRNA-Seq data using the Louvain community detection algorithm. This generated a total of n=16 clusters. We next annotated the clusters with cell types using singleR algorithm and well-established cell type markers. We identified monocyte, macrophage, T-cell and tumor cells (Figure 4A). We next identified large scale CNV events using CaSpER^42^ to elucidate the effect of CNVs on velocity (Supplementary Figure 1). We noticed the tumor cluster 0 and 8 not harboring chr11p and chr18q deletion. We believe that these cells represent less aggressive cell clones within the tumor compared to other tumor cell clusters. We supported our hypothesis by calculating an aggressiveness score for each cell using gene signatures of aggressive and non-aggressive tumors identified from our previously published bulk RNA-Seq expression data^43^. We have observed that cluster 0 and cluster 8 got a higher score for non-aggressive meningioma tumors. We next inferred RNA velocity in our scRNA-Seq meningioma data and projected the velocities on to our previously defined UMAP embeddings. We observed a similar finding that non-aggressive meningioma cells are moving towards aggressive meningioma cells in velocity based trajectory analysis (Figure 4B).

**Figure 4.**
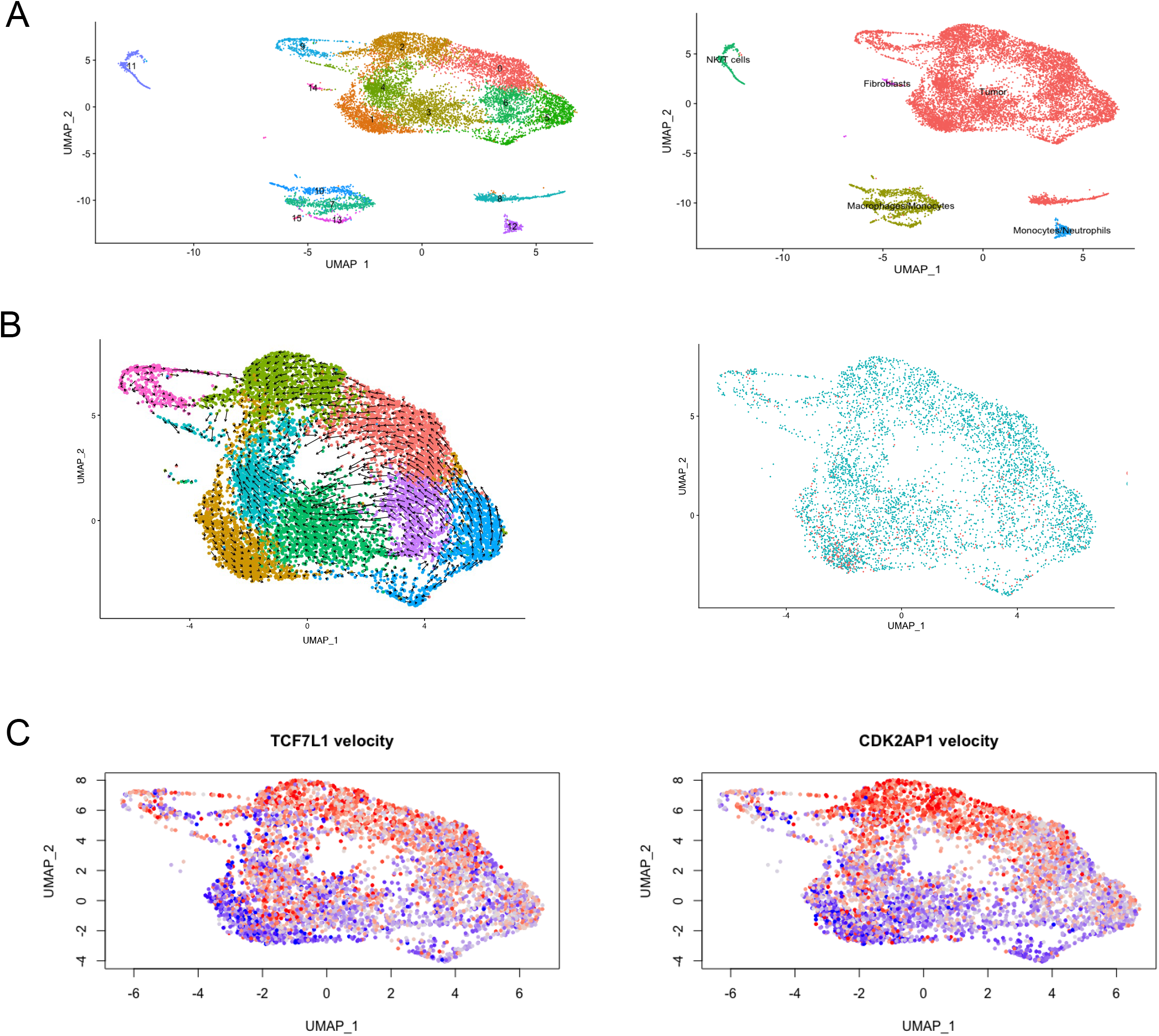
**A)** scRNA-Seq meningioma clustering and annotation **B)** trajectory of scRNA-Seq meningioma data using RNA Velocity and aggressiveness score for each cell using gene signatures of aggressive and non-aggressive tumors **C)** *TCF7L1* which is a mediator of the Wnt signaling pathway, and *CDK2AP1* which epigenetically regulates embryonic stem cell differentiation, have increased transcriptional dynamics in more aggressive tumor cells.

We next applied our scRegulocity algorithm on our single cell meningioma data. We observed that hypoxia related TFs such as *DDIT3 and NR3C1* have repressed transcriptional dynamics in less aggressive tumor cells. Similarly, *TCF7L1* which is a mediator of the Wnt signaling pathway^44^, and *CDK2AP1* which epigenetically regulates embryonic stem cell differentiation^45^, have increased transcriptional dynamics in more aggressive tumor cells (Figure 4C).

## Methods

### Intron/Exon Read Quantification

The velocity estimation starts by quantifying the intron/exon read counts. The basic idea is that genes that exhibit increase (decrease) in expression will harbor more (less) reads on the introns compared to the baseline exonic reads. This basic idea is used to estimate and assign expression velocity estimates to each gene. scRegulocity contains a module that counts intronic (unspliced) and exonic (spliced) read counts for each gene in each sample. and normalize the counts using total number of reads in each sample. scRegulocity has a specific module to perform read quantifications in an integrated manner so that there is no dependence on the other packages. Specifically, scRegulocity makes use of the “CB:Z” tags in the reads to first assign each read to a cell then identified whether the read belong to an intron, exon, or an intron-exon junction, i.e., unspliced reads. scRegulocity keeps track of 3 different counters 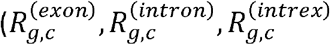, corresponding to exonic, intronic and boundary read counts for cell at index *c*, and the gene at index *g*) for each gene *g* and concurrently keeps track of these counts while quantification is being performed. After quantification is finished, the count matrix (genes in the rows, cells in the columns) is saved in a tab-delimited file. The C++ code for quantification, can be downloaded from GitHub at https://github.com/harmancilab/IntrExtract/. Our method takes a bam file as an input and outputs 3 matrices where the rows are the genes and the columns are exons, introns or ambigious reads.

### Velocity Estimation

scRegulocity includes a flexible and integrated velocity estimation module that can be parametrized by the user. The velocity estimation takes the intron/exon read counts matrix as input. The number of reads mapping to the introns and intron-exon boundaries are generally 1-2 orders of magnitude smaller than that of reads mapping to the exons. This is expected since exonic reads dominate the transcripts that are sequenced in RNA-seq protocols. For velocity estimation, it is necessary to obtain a robust estimate of the ratio of reads that are mapping on introns (and intron/exon boundaries) and the exonic reads, which is proportional to the expression velocity. To provide an estimate of this ratio, scRegulocity performs a linear regression between the unspliced 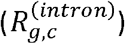 and spliced read counts of all genes 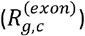, as it is currently a well-established approach to estimate velocity ^28,29^:

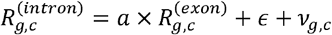

where intronic read counts are modeled as an ordinary linear model of the exonic read counts, *a* indicates the slope, *ϵ* represents a random noise term to include technical noise, and *v*(*g*) represents the velocity of the expression. From this model, the general linear trend between unspliced and spliced reads quantifies a gene-specific spliced/unspliced read counts. This effect represents mostly a technical component (see above) whereby the highly expressed genes will contain more reads on the introns and is removed by subtracting the linear component from the spliced/unspliced ratios of the genes. The residual intronic (unspliced) read counts are used as the final velocity estimates for all genes. To make the estimate more robust, the linear model uses extremes of the spliced/unspliced ratios, specifically the upper and lower quantiles, which is set to %*q*_*v*_ = 0.05 by default, an approach similar to the velocyto workflow ^46^. This parameter can be changed to make the linear trend removal more stringent or more relaxed. To test for other factors that may bias velocity estimates, we have tested the model by including covariates such as read-mappability and GC content. We observed that these covariates do not significantly improve velocity estimates and therefore are by default not explicitly included in the velocity estimation module of scRegulocity. The final velocity estimation step will yield a matrix of RNA velocity estimates where each row represents a gene and column represents cells.

### Building the Cell-Cell Gradient Graph

scRegulocity identifies the velocity switching patterns using a graph-based approach. The target is to identify two sets of cells that are neighboring in the embedding such that there is a coordinated switch in the expression velocities of all cells from one set of cells to the other set of cells. In other words, we would like to identify a strong coordinated gradient between two sets of neighboring cells such that the velocities are switched between the two sets of cells. First, the embedding coordinates of the cells are analyzed and a pairwise cell-cell distance matrix is generated. Given a K-dimensional embedding, this can be simply computed by:

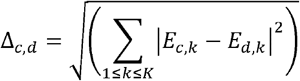

Where *E*_*c,k*_ denotes the embedding coordinates of cell *c* at coordinate *k*. Based on the cell-cell distance matrix, scRegulocity uses a neighborhood parameter *σ*_*v*_ that denotes the largest radius at which cells are deemed as neighbors of each other. For each *σ*_*v*_ *-*neighborhood, scRegulocity forms a graph where the nodes are placed on cells and edges are placed between cells that are in each other’s -neighborhoods. For each edge, we assign a weight based on the absolute value of velocity difference between the cells that are connected by the edge. Given the velocity estimates for the gene *g*,

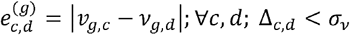

where 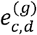 represents the weight of the edge that connects cells *c* and *d*, which are in the *σ*_*v*_ -neighborhood of each other. The edges are also assigned directions based on the sign of difference between the velocities of the cells. In this representation, the edges represent discrete units of velocity gradients that will be used to detect concordant velocity-switches. However, we observed that most of the edges do not provide useful information as they represent random and weak gradient vectors between cells. In addition, processing of the cell-cell network with all the edges increases computational cost. To overcome this, the edges are pruned with respect to the weight threshold, *e*_*min*_, weights so that the weak gradients are excluded from analysis, i.e., for a gene *g*, the edges 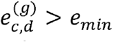 are retained from edge filtering. Currently, *e*_*min*_ *=* 2 is used by default as a stringent weight threshold that also provides enough power to identify velocity switching patterns. After the edges are filtered, the weakly connected components of the graph (all cells are connected to each other without regard to the direction of edges) are identified using breadth-depth first search algorithm^47^. We refer to these components as candidate velocity switching subnetworks because they contain the sets of cells where the velocity switching events take place. For a gene *g*, Each candidate is defined by the connected cell-cell edge subnetwork with the filtered edges:

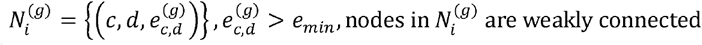

where 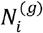 denotes *i* ^*th*^ subnetwork for gene *g*.

### Detection of Velocity-Switching Patterns

The cell-cell gradient information in each network is expected to correspond to one velocity switching pattern. In order to detect abrupt changes in velocity, the candidate subnetworks are tested to identify whether there is a significant velocity switching pattern in them. The first test checks for concordance of the gradient in the subnetwork. For this, scRegulocity computes the directions of the gradients defined by each edge. The orientation of the gradient vectors are assigned using the coordinates of the cells that they are connecting. Given the 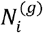 the list of all edges in the *i* ^*th*^ cell-cell subnetwork, the direction and weights of all the edges in the subnetwork are extracted:

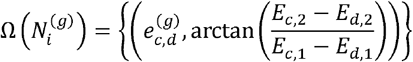

where 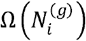 denotes pairs of edge we ientation angle, which is computed using inverse tangent (i.e., arctan) of the height/width ratio of the rectangle formed on the embedding coordinates of the cells at the ends of edge. scRegulocity also uses the orientation and weights of the edge to compute the aggregated gradient vector, which is used in visualization:

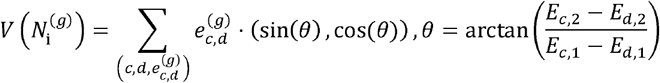

This vector is a 2-element vector of embedding coordinates that is vector summation of the unit vectors along each gradient vector after the unit vectors are multiplied by the weight of the corresponding edge. 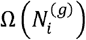 contains the direction and strength information of the cell-cell velocity gradients. scRegulocity uses this information to statistically test whether there is an enrichment of high weighted cell-cell gradients along the same orientation. For this, Rayleigh test, which performs a statistical test of whether the sample of orientations are different from a random distribution that are sampled uniformly over the unit circle. Rayleigh test also considers the weights naturally to weigh each orientation so that the gradient vectors with higher weights contribute more on the test statistic. For a subnetwork 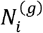, we input 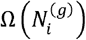 into the Rayleigh test to assess the significance of whether the weighted edges are distributed uniformly. The subnetworks are filtered with respect to the p-value threshold. As a further test of the significance of a gradient pattern on the subnetwork, scRegulocity computes the spatial correlation using moranI test on RNA velocity gradient vectors between the cells near the velocity switch. This test takes as input the embedding coordinates of the cells and the velocity values on each cell, i.e., *v*_*g,c*_. The significance of the pattern on the embedding are computed with respect to a null model where the velocities are randomly distributed.

scRegulocity uses the p-value of the moranl test to filter out subnetworks that do not exhibit significant spatial patterns. After this step, the subnetworks are scored and filtered and we have the final set of velocity switching patterns at the cells that are contained in the filtered subnetworks. For each gene, a number of subnetworks are identified.

### Clustering of genes with respect to velocity switch patterns

We cluster genes based on velocity switch patterns. For this scRegulocity build a velocity direction matrix for each subnetwork for each gene. The matrix entries contain the positive (+1) and negative (−1) values indicating the direction of velocity in all cells. The matrix contains genes in the rows and cells in the columns. The genes are clustered with respect to similarity of the (discretized) velocity direction values using Euclidean distance matrix with partitioning around medoids clustering method^48^. We hypothesize that the genes that share the velocity switch pattern on same cells exhibit similar abrupt coordinated changes in the gene regulatory processes. To uncover this, scRegulocity performs pathway enrichment analysis on the set of genes within each cluster using enrichR R package^49^.

### Transcription Factor (TF)-Target Velocity Regulation

For each TF gene with at least one significant velocity switching pattern, scRegulocity evaluates the targets (via motif and regulatory-network databases ^50^. Next, the expression levels and velocities of the targets are checked for correlation with the regulator’s expression levels and velocities. These are visualized in a network view at the velocity and expression level.

### Visualization

scRegulocity contains visualization modules that automatically take the outputs of the algorithm as input and generate visualizations of (1) the significant velocity switching patterns with a directional distribution of gradients on subnetworks, (2) gene-clusterings, and (3) TF-target networks are output and visualized by scRegulocity so that the results can be easily interpreted and assessed in terms of biological significance with respect to the tested hypotheses.

## Discussion

We present an algorithm, scRegulocity, for identification and visualization of driver genes and regulatory networks within transient cell states and types. We demonstrated that scRegulocity can deconvolute the transcriptional drivers using RNA Velocity with our graph based algorithm. We present several examples where scRegulocity effectively complements the existing set of RNA Velocity analysis tools and gives insight into the understanding of cell-state transitions in diverse systems. ScRegulocity can extend the utility of RNA-seq datasets beyond just transcriptional profiling.

In conclusion, scRegulocity is a method that generates RNA Velocity from single cell RNA-seq data and infers driver transcription factors and transcriptional modules to guide the discovery and understanding of the cellular states. Our results show that scRegulocity can more accurately recover dynamic transcription factor (TF) modules compared to whole transcriptome single cell expression RNA data.

## Acknowledgement

This study is partially funded by the NIH grant R01CA241930.

